# Senescence of cortical neurons following persistent DNA double-strand breaks induces cerebrovascular lesions

**DOI:** 10.1101/2023.01.26.525738

**Authors:** Caroline A. Kopsidas, Clara C. Lowe, Jun Zhang, Wenjun Kang, Xiaoming Zhou, Yuanyi Feng

## Abstract

DNA double strand breaks (DSBs), neuroinflammation, and vascular alterations in the brain are all associated with neurodegenerative disorders. However, the interconnections between these neuropathological changes and how they act synergistically to promote irreversible neurodegeneration remain unclear. Here we show that abrogating the BRCA1-associated protein Brap in cerebral cortical neurons, as opposed to vascular endothelium cells, causes cerebrovascular defects. This non-cell autonomous effect is mediated by cellular senescence resulting from persistent neuronal DSBs. We show that in the state of senescence, there is a massive upregulation of genes involved in cell secretion, inflammatory responses, and vascular changes, which coincides with cerebral microclots and microbleeds. The vascular lesions intertwine with neuroinflammation and exacerbate neuronal DSBs, culminating in oxidative stress, metabolic alteration, and downregulation of genes essential for neuronal function. By demonstrating the cerebrovascular impact of cortical neuronal DSBs, our data suggest that senescence-associated secretory phenotype can initiate brain-wide neurodegeneration.

## Introduction

Genome instability with increased DNA double strand breaks (DSBs) is detrimental to neurons of the brain and has been implicated in diverse neurodegenerative disorders (NDs)^1–6^. However, the etiology of NDs is multifaceted, the progressive decline in number and activity of neurons exhibits significant comorbidity with glial and cerebrovascular abnormalities^7–10^. Thus, determining the mechanism by which neuronal DSBs interact with glial and vascular dysfunction in the brain is essential for understanding the pathophysiology that underlies brain-tissue-wide degeneration.

By studying chromatin remodeling in neurogenesis, we have recently shown that DSBs in neocortical neurons may arise from compromised epigenetic silencing of pericentric satellite DNA in mice with genetic deficiencies of Nde1 (neurodevelopment protein 1) and its interacting Brap (BRCA1-associated protein)^11^. The long tandem repeats of satellite DNA are intrinsically unstable and refractory to repair^12^. We found that the de-repression of satellite repeats in Brap neural progenitor cell (NPC) or neuronal conditional knockout (cKO) mice (Brap^cKONPC^ and Brap^cKONeuron^, respectively) results in persistent DSBs in cortical neurons and subsequently a senescence state along with accelerated neurodegeneration and shortened lifespan^13^. In vivo, cellular senescence is characterized by the distinctive senescence associated secretory phenotype (SASP)^14–20^, through which it imposes micro-environmental effects to the nearby tissue. By secreting a complex mixture of pro-inflammatory and pro-oxidative cytokines, chemokines, growth factors, and proteases, SASP can induce inflammation and aging^21–26^. Indeed, cellular senescence in cerebral cortices of Brap^cKONPC^ mice was accompanied by the upregulation of secreted molecules with roles in not only immune regulation but also neural networking and cerebrovascular activities^13^, which suggests that senescence is a mediator by which neurons with DSBs impair their neighboring cells in the surrounding brain tissue.

In this study, we provide proof of concept evidence for the non-cell autonomous effect of neuronal DSBs. Through studying pathological consequences of mice in which Brap is abrogated in various cell types of the brain, we report the finding that Brap loss of function (LOF) in cortical excitatory neurons can cause diverse cerebrovascular lesions. These vascular damages co-occur with neuroinflammation and synergize with DSBs, resulting in brain metabolic and redox dysfunction to accelerate neurodegeneration. These results suggest that SASP is a mechanism through which genotoxic insults, especially DSBs in repetitive sequences of cortical neurons, impinge on broad brain structures and regions.

## Results

### Brap abrogation in cortical neurons causes diverse cerebrovascular lesions

While Brap LOF results in DSBs and cellular senescence of replication competent cells in culture, DSBs in the cerebral cortex were specifically exhibited in neurons of mice in which Brap was abrogated in NPCs that give rise to cortical glutamatergic neurons, astroglia, and oligodendrocytes by Emx1-Cre^27^ or in postnatal cortical and hippocampal neurons by Thy1-Cre^28^ (referred to as Brap^cKONPC^ and Brap^cKONeuron^, respectively, or collectively as neural Brap^cKO^ mice hereafter unless noted specifically)^13^. These mutant mice show rapid brain degeneration and premature death starting at the young adult age (Figure 1A). To understand the cause of mortality and neurodegeneration, we monitored neural Brap^cKO^ mice closely and found death in more than half of them occurred suddenly without any sign of fatal conditions the day before, though the mutant mice were overtly hyperactive and some showed transient episodes of seizures (Supplemental videos 1 and 2). We were able to catch about 40 male and female neural Brap^cKO^ mice that showed sudden onset of fatal symptoms, including extreme lethargy, labored breathing, or hemi/monoplegia, which suggested death was imminent (Figure 1B, Supplemental videos 3 and 4). On examination, about 20% of them showed visible subarachnoid and/or intracerebral hemorrhage (Figure 1C). Brain histological and immunohistological analyses of neural Brap^cKO^ mice revealed widespread cerebral microvascular lesions (Figure 1D, E). While some intracerebral blood vessels appeared leaky with albumin extravasation, it frequently co-occurred with vascular occlusions in these mutant brains. These data suggest that the sudden death of neural Brap^cKO^ mice was due to both ischemic and hemorrhagic damages of cerebral blood vessels.

**Figure 1.**
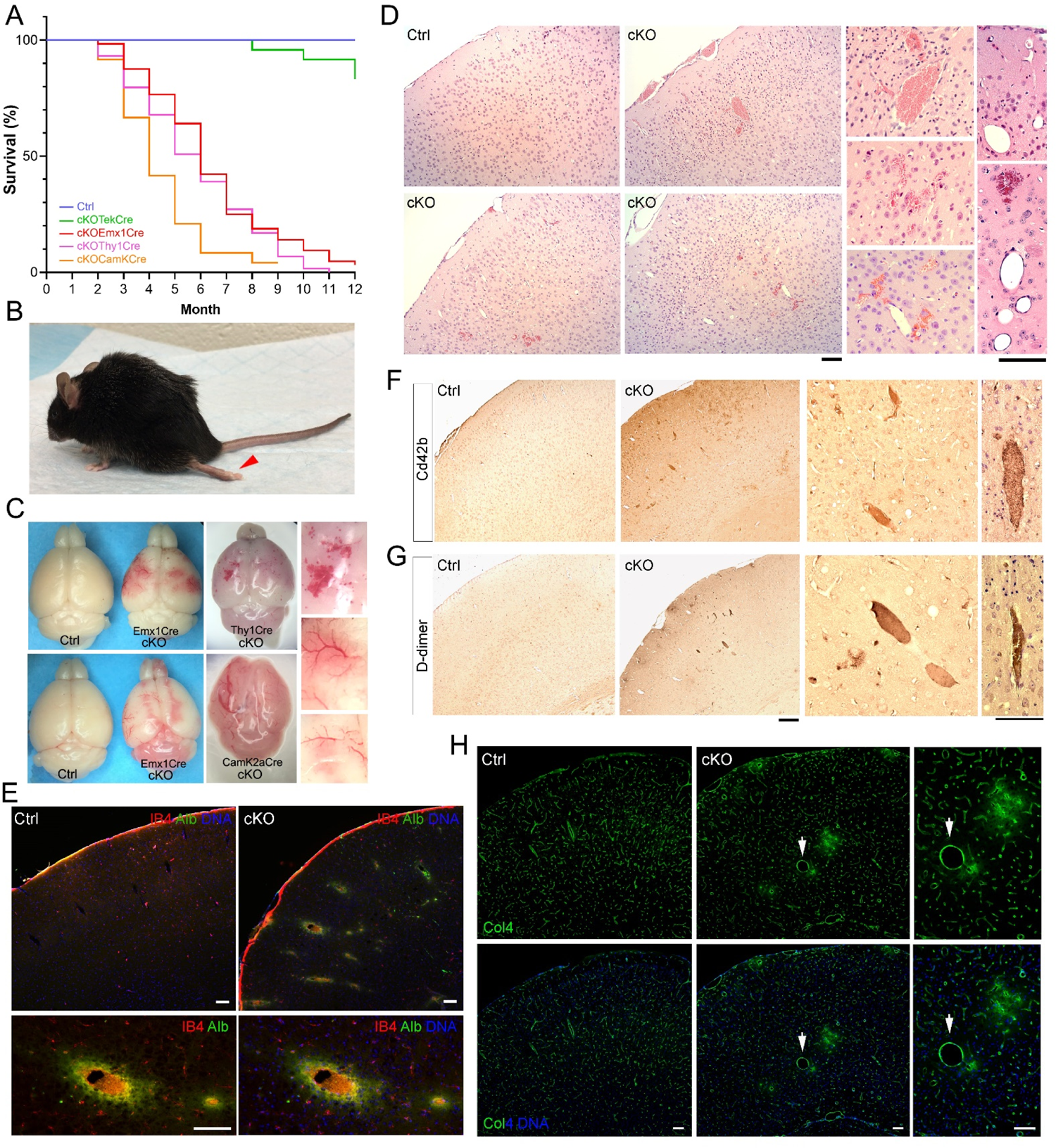
Brap loss of function in cortical neurons results in midlife mortality and diverse cerebrovascular lesions. (A) Kaplan Meier curves of mice with Brap conditional knockout in vascular endothelial cells (Tek-Cre, n=24), dorsal telencephalon neural progenitor cells (Emx1Cre, n=65), postnatal cortical and hippocampal neurons (Thy1Cre, n=60), excitatory neurons (Camk2aCre, n=25), or their control counterparts (n=47). (B) A neural Brap^cKO^ mouse that showed sudden paralysis of a hindlimb (monoplegia). The arrow indicates the dragging hindlimb. See also supplemental video. (C) Representative brain images of neural Brap^cKO^ mice that exhibited subarachnoid hemorrhage. (D) H&E histology images show diverse cerebrovascular lesions, including subarachnoid and intracerebral hemorrhage, as well as microbleeds, aneurysm, and occlusion of small cerebral blood vessels in neural Brap^cKO^ mice. (E) Fluorescence immunohistological images of cortical sections double stained by Isolectin 4 (IB4, in red) and an albumin antibody (Alb, in green). Note the co-existence of cerebral vascular occlusion and albumin extravasation in neural Brap^cKO^ mice. Nuclear DNA was stained with Hoechst 33342. (F&G) Representative immunohistochemistry images of cortical sections stained by antibodies against Cd42b or D-dimer. (H) Representative fluorescence immunohistological images of cortical sections stained by an antibody to collagen IV (Col4, in green). The arrow indicates a bulging cerebral blood vessel in neural Brap^cKO^ mice. Nuclear DNA was stained with Hoechst 33342. Bars: 100um.

To verify the co-occurrence of cerebrovascular micro clots and microbleeds, we examined over 50 neural Brap^cKO^ mice between weaning and 12 months of age. About ∼60% of the neural Brap^cKO^ mice showed cerebrovascular occlusions enriched with Cd42b, a marker for platelet activation and clot formation (Figure 1F). The presence of cerebrovascular clots or thrombosis was further confirmed by high levels of D-dimer, the product of fibrin degradation (Figure 1G). In addition to blood clots, cerebral aneurysm, defined by isolated blood vessel dilation, balloons, or bulges, was also shown by about 20% of the neural Brap^cKO^ mice (Figure 1D, H). These vascular lesions affected predominantly small cerebral vessels of < 50um in diameter, distributed sporadically throughout the cerebrum in the subarachnoid space, the neocortex, the white matter, and sometimes in other brain regions (Figure 1D-H, Figure S1A). The cerebral vascular defects were comparable in Brap^cKONPC^ and Brap^cKONeuron^ mice but were not shown by their control counterparts (Brap^flox/wt^; Cre-), which suggests that Brap LOF in the cortical parenchyma, can cause prevalent cerebral small vessel lesions.

To ascertain that the cerebrovascular defects were caused by Brap LOF in neurons, we generated and analyzed mice in which Brap was deleted in excitatory neurons of the forebrain and the CA1 of hippocampus by Camk2a-Cre^29^ (Brap^cKOENeuron^) and vascular endothelial cells by Tek-Cre^30^ (Brap^cKOEC^), respectively. In contrast to the mild survival impairment and subtle brain defects of Brap^cKOEC^ mice, Brap^cKOENeuron^ mice, in which Brap was broadly abrogated in excitatory neurons of the neocortex, hippocampus, striatum, and other brain structures, showed not only cerebrovascular defects but also a stronger survival disadvantage than Brap^cKONPC^ and Brap^cKONeuron^ mice (Figure 1A, Figure S1B, C). Therefore, the shared phenotype of mice with loss of Brap commonly in cortical excitatory neurons provides compelling evidence for a non-cell autonomous neuronal impact on cerebral blood vessels. This finding fully agrees with the notion that persistent DSBs resulting from Brap LOF can drive cortical neurons to a state of senescence and affect cells in the nearby cortical tissue by SASP.

### Co-upregulation of inflammatory, coagulation, hypoxia, ROS, and DNA damage responses

To determine the molecular profile by which neuronal Brap LOF affects the cerebral vasculature, we performed transcriptome analysis of Brap^cKONPC^, Brap^cKONeuron^, and Brap^cKOEC^ mice at 3-4 months of age along with age matched control mice (Brap^flox/wt;^ Cre-). We analyzed poly(A) enriched cortical mRNA and obtained more reads than the previous RNAseq of total cortical RNA from Brap^cKONPC^ mice^13^. Compared to control mice, Brap^cKONPC^ and Brap^cKONeuron^ mice showed a total of 3,301 and 1,962 differentially expressed genes (DEGs), respectively, whereas Brap^cKOEC^ mice only showed 13 DEGs (FDR<0.05; Fold Change >1.5) (Figure 2A). Gene ontology (GO) analyses revealed that the 1,570 and 1,205 significantly upregulated DEGs in Brap^cKONPC^ and Brap^cKONeuron^ mice were enriched most significantly in cellular components associated with secretory pathways, including cytoplasmic vesicles, extracellular exosomes and vesicles, secretory granules and vesicles, as well as the cell substrate junction (Figure 2B). Such remarkable increase in the expression of genes involved in cell secretion strongly supports the non-cell autonomous effects of SASP by Brap deficient neurons.

**Figure 2.**
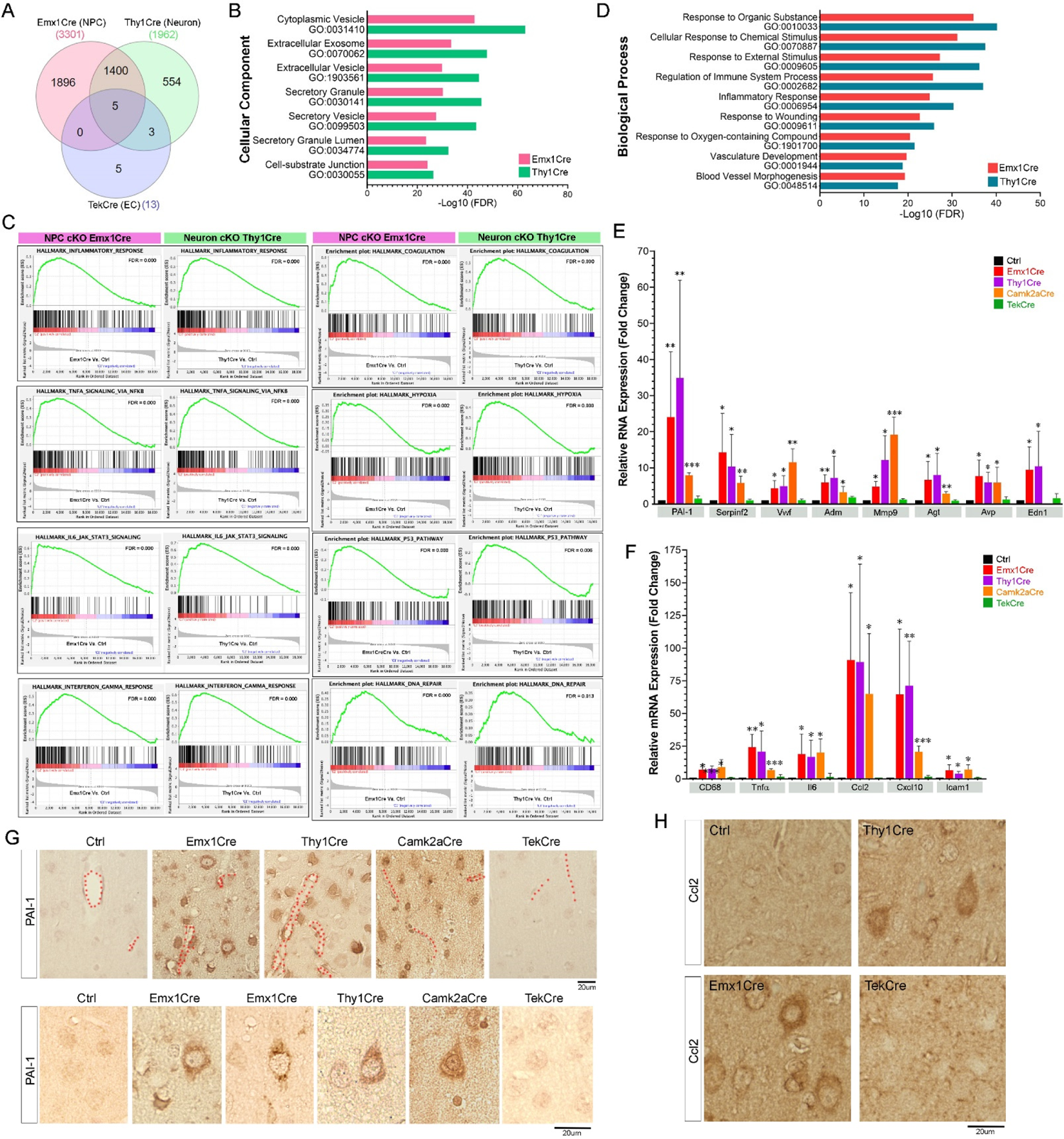
Significant upregulation of genes mediating cellular secretion, inflammation, vascular changes, and DNA damage responses in cortical tissues of neural Brap^cKO^ mice. (A) Venn diagram of differentially expressed genes in cerebral cortices of mice with Brap conditional knockout in vascular endothelial cells (Tek-Cre, n=5), dorsal telencephalon neural progenitor cells (Emx1Cre, n= 6), or postnatal cortical and hippocampal neurons (Thy1Cre, n=5) at 3-4 months of age relative to age-matched control mice (n=4). (B) Gene Oncology Cellular Component (CC) terms identified in significantly upregulated cortical protein-coding genes of mice in which Brap was conditionally abrogated in dorsal telencephalon neural progenitor cells (Emx1Cre), or postnatal cortical and hippocampal neurons (Thy1Cre). Shown are selected enrichment terms and corresponding FDR values. (C) Selected GSEA of hallmark pathways enriched in unique cortical genes identified from mice with Brap conditional knockout in dorsal telencephalon neural progenitor cells (Emx1Cre), or postnatal cortical and hippocampal neurons (Thy1Cre). Shown are enrichment score plots with FDR values compared to cortical genes identified from control mice. (D) Gene Oncology Biological Processes (BP) terms identified in significantly upregulated cortical protein-coding genes of mice in which Brap was conditionally abrogated in dorsal telencephalon neural progenitor cells (Emx1Cre), or postnatal cortical and hippocampal neurons (Thy1Cre). (E) RT-qPCR analyses of selected vasoactive genes in the cortical tissue of mice with Brap conditional knockout by Emx1Cre, Thy1Cre, Camk2aCre, or Tek-Cre mice at 3 months of age. Shown are relative levels of mRNA expression compared to that of age-matched control mice. Error bars represent SD; *p < 0.05; ** p< 0.01; *** p< 0.001, by ANOVA tests. (F) RT-qPCR analyses of selected inflammatory genes in the cortical tissue of mice with Brap conditional knockout by Emx1Cre, Thy1Cre, Camk2aCre, or Tek-Cre mice at 3 months of age. Shown are relative levels of mRNA expression compared to that of age-matched control mice. Error bars represent SD; *p < 0.05; ** p< 0.01; *** p< 0.001, by ANOVA tests. (G) Representative immunohistochemistry images of cortical sections stained by antibodies against PAI-1. Note the high level of PAI-1 in cortical neurons of neural Brap^cKO^ mice. Cerebral blood vessels are marked by doted lines. (H) Representative immunohistochemistry images of cortical sections stained by antibodies against Ccl2. Note the high level of Ccl2 in cortical neurons of neural Brap^cKO^ mice.

We next performed gene set enrichment analysis (GSEA) to compare the transcriptome of Brap^cKONPC^ or Brap^cKONeuron^ mice relative to their control counterparts. These transcriptome data demonstrated the phenocopy of Brap LOF in cortical NPCs and neurons with respect to significant enrichment of genes implicated in over 20 hallmark pathways, including inflammatory response, TNFα signaling via NFkB, IL6 JAK STAT3 signaling, interferon-γ response, coagulation, hypoxia, P53 pathway, and DNA repair (Figure 2C, Figure S2A). Whereas very few hallmark pathways were significantly over-represented by mRNAs in control relative to Brap^cKONPC^, and Brap^cKONeuron^ cortices (Figure S2B). Likewise, GO analyses of significantly upregulated DEGs in Brap^cKONPC^, and Brap^cKONeuron^ cortices revealed over-representation of genes associated with biological processes in inflammatory or vascular responses to stimuli, wound, and oxygen-containing compounds (Figure 2D). Collectively, our data suggest that, in spite of the activation of p53 pathway and DNA repair, DSBs in Brap deficient cortical neurons persist and drive them to a senescent state, under which upregulated expression and secretion of inflammatory and vasoactive molecules lead to multiple secondary effects on local astroglia, microglia, and blood vessels.

We confirmed increased mRNA of several vasoactive molecules in neural Brap^cKO^ cortices by RT-qPCR. These include PAI-1(Serpine1), a bona fide SASP molecule that inhibits tissue plasminogen activator (tPA) and blocks clot breakdown^31–33^, α2-antiplasmin (Serpinf2), a potent plasmin inhibitor that stops fibrinolysis^34–36^, Vwf, a key player in thrombus formation by facilitating platelet adhesion to damaged vascular walls^37^, Adm (Adrenomedullin), a vasodilator peptide^38, 39^, Mmp9, a matrix metalloprotease that acts on the vascular wall^40–42^, Agt (Angiotensinogen), the precursor for all angiotensin peptides that promote vasoconstriction and blood pressure increase^43^, Avp (Vasopressin), a peptide hormone that constricts arterioles and raises blood pressure^44, 45^, and Edn1 (endothelin-1), a strong and long-acting vasoconstrictive peptide that can induce stroke^46–48^(Figure 2E). These factors can act differentially and/or synergistically to induce heterogeneous vascular changes. Their significant upregulation in the cortical tissue indicates the loss of neurovascular hemostasis, which explains the diverse ischemic and hemorrhagic defects in cerebral small vessels and the sudden midlife mortality of neural Brap^cKO^ mice.

All neural Brap^cKO^ mice with significant upregulation of vasoactive molecules also showed increased expression of genes associated with neuroinflammation and microglia activation, such as Tnfα, Il6, Ccl2, Cxcl10, Icam1, and CD68 (Figure 2F). As inflammation is a major hallmark of cellular senescence, these data are fully in line with SASP-mediated dual-effect of intractable neuronal DSBs on both inflammatory and vascular cells.

We further confirmed that the strongly elevated PAI-1 and Ccl2 proteins in the cortical tissue of neural Brap^cKO^ mice were indeed produced by senescent neurons. We showed that both molecules were highly enriched in some cortical neurons of neural Brap^cKO^ mice but largely absent in those of the Brap^cKOEC^ and control brains (Figure 2G, H). Therefore, by producing vasoactive and inflammatory molecules, neuronal senescence can cause brain tissue-wide vascular defects and neuroinflammation.

### Intermingle of cerebrovascular lesions with neuroinflammation and neuronal DSBs

To further reveal the paracrine effect of neuronal senescence, we accessed the spatial correlation of cerebral vascular lesions with inflammatory responses of glial cells and neuronal DSBs in neural Brap^cKO^ mice. While astrogliosis and microgliosis are highly elevated in Brap^cKONPC^ and Brap^cKONeuron^ cortices^13^, we found the activated astrocytes and microglia were spatially associated with vascular lesions. We detected reactive astrocytes by Gfap immunoreactivity and active microglia by their strong Iba immunosignals and enlarged roundish cell body with retracted ramification, and found they were highly abundant in the vicinity of occluded, bulged, or ruptured cerebral blood vessels in neural Brap^cKO^ mice (Figure 3A-D; Figure S3A-C). Furthermore, we showed that damaged and misshapen cerebral vessels were also surrounded by neurons with high levels of γH2A.X, a biomarker for DSBs (Figure 3E, F; Figure S3D). Although vascular injury can trigger gliosis, and conversely, neuroinflammation may augment vascular defects, their co-occurrence with neuronal DSBs suggests they were both initiated by SASP.

**Figure 3.**
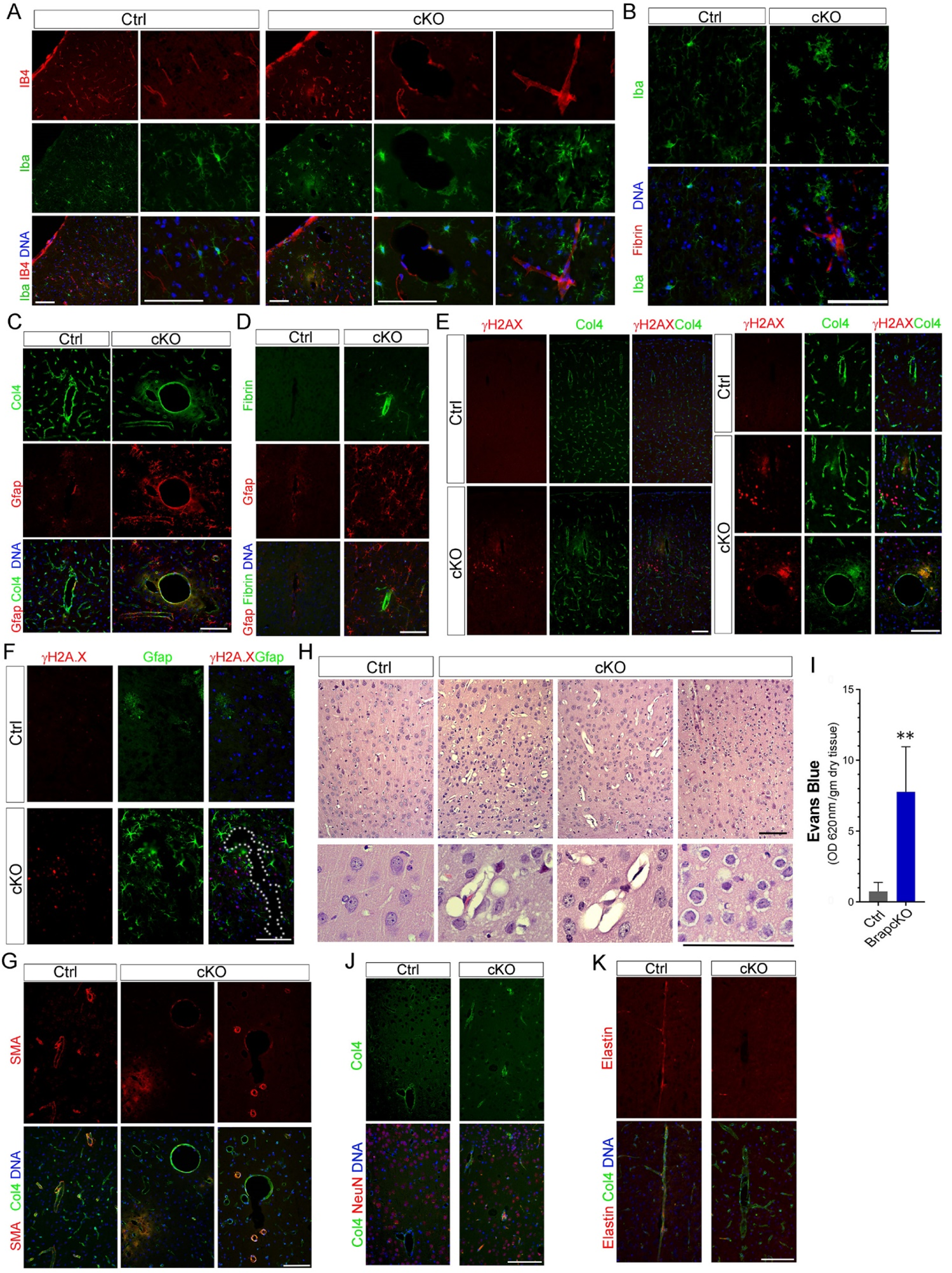
Spatial correlation of DSBs with intertwined cerebral cortical glial activation and vascular lesions in neural BrapcKO mice. (A) Representative fluorescence immunohistological images of cortical sections stained by Isolectin 4 (IB4, in red) and the microglia marker Iba (in green). Note the enlarged microglia surrounding microbleeds and bulging cerebral blood vessels in neural Brap^cKO^ mice. (B) Representative fluorescence immunohistological images of cortical sections double stained by antibodies to fibrin (in red) and Iba (in green). Note the enlarged microglia next to a blood vessel with fibrin occlusion in the neural Brap^cKO^ cortex. (C) Representative fluorescence immunohistological images of cortical sections double stained by antibodies to Gfap (in red) and Col4 (in green). Note the reactive astrocytes with high Gfap surrounding a bulging cerebral blood vessel in the neural Brap^cKO^ cortex. (D) Representative fluorescence immunohistological images of cortical sections double stained by antibodies to Gfap (in red) and fibrin (in green). Note the strong Gfap in the region with vascular fibrin occlusion in the neural Brap^cKO^ cortex. (E) Representative fluorescence immunohistological images of cortical sections double stained by antibodies to the DSB marker γH2A.X and Col4, showing high abundance of γH2A.X signals nearby microbleeds and bulging blood vessels in the neural Brap^cKO^ cortex. (F) Representative fluorescence immunohistological images of cortical sections double stained by antibodies to the DSB marker γH2A.X (in red) and activated astrocytes Gfap (in green), showing the high abundance of both γH2A.X and Gfap signals in the vicinity of a misshapen blood vessel (marked by dotted outlines) in the neural Brap^cKO^ cortex. (G) Representative fluorescence immunohistological images of cortical sections double stained by antibodies to smooth muscle actin (SMA, in red) and Col4 (in green), showing reduced SMA in the vascular wall of neural Brap^cKO^ cortex. (H) H&E histology images show cerebral edema and enlarged perivascular space in neural Brap^cKO^ mice. (I) Evans blue assay reveals increased blood brain barrier permeability in pre-symptomatic Brap^cKONPC^ mice. Error bars are SD. ** p< 0.01 (J&K) Representative fluorescence immunohistological images of cortical sections double stained by an antibody to Col4 (in green) and antibodies to NeuN or elastin (in red), showing the accumulation of Col4 with reduced elastin in the cerebral blood vessels of surviving neural Brap^cKO^ mice. Nuclear DNA was stained with Hoechst 33342 in blue. Bars: 100um.

We also detected structural defects in cerebral blood vessels of neural Brap^cKO^ mice. Damaged vascular wall, evidenced by loss of smooth muscle actin, was shown by those cerebral vessels with occlusions, aneurysm or ruptures in the acute stage of vascular lesions (Figure 3G, Figure S3E). In addition, common signs of intracerebral small vessels defects, such as cerebral edema with vacuolar appearance of the gray matter and enlargement of perivascular space, were presented by neural Brap^cKO^ brains (Figure 3H). Furthermore, we detected a significant increase in blood brain barrier (BBB) permeability in presymptomatic Brap^cKONPC^ mice (Figure 3I). In those Brap^cKONPC^ mice that survived past 6 months, there were elevations of collagen deposition and loss of elastin in cerebral blood vessels (Figure 3J, K), which are signs of basement membrane thickening and increased vascular stiffness. These vascular changes suggest chronic cerebral hypoperfusion in the surviving neural Brap^cKO^ mice.

### Brain metabolic alteration with increased ROS, neuronal necrosis, and down-regulation of genes essential for neuron function

We next sought to determine the impact of vascular lesions on brain metabolism and neural function in neural Brap^cKO^ mice. The brain relies on uninterrupted supply of glucose and oxygen from blood flow to support neuronal activities. Glucose in the brain is mainly used to generate ATP through glycolysis and oxidative phosphorylation (OXPHOS) in the tricarboxylic acid (TCA) cycle, during which NAD^+^ (nicotinamide adenine dinucleotide) is reduced to NADH to donate electrons in the respiratory chain. Alternatively, glucose in the brain can be processed by the pentose phosphate pathway (PPP) to generate NADPH (reduced form of nicotinamide adenine dinucleotide phosphate, NADP^+^) and ribose 5 phosphate, which is essential for diverse biosynthesis and reactive oxygen species (ROS) clearance (Figure 4A). Corroborated with compromised vascular circulation, the cortical tissue of neural Brap^cKO^ mice showed a significant decrease in the level of NADH but increases in levels of NADPH and ROS (Figure 4B-D). This suggested that vascular defects not only led to glucose and oxygen deprivation but also a glucose metabolic reprograming: instead of being broken down by glycolysis and OXPHOS to generate ATP, glucose in the neural Brap^cKO^ brain was shunted to PPP, producing NADPH to scavenge ROS and combat oxidative stress.

**Figure 4.**
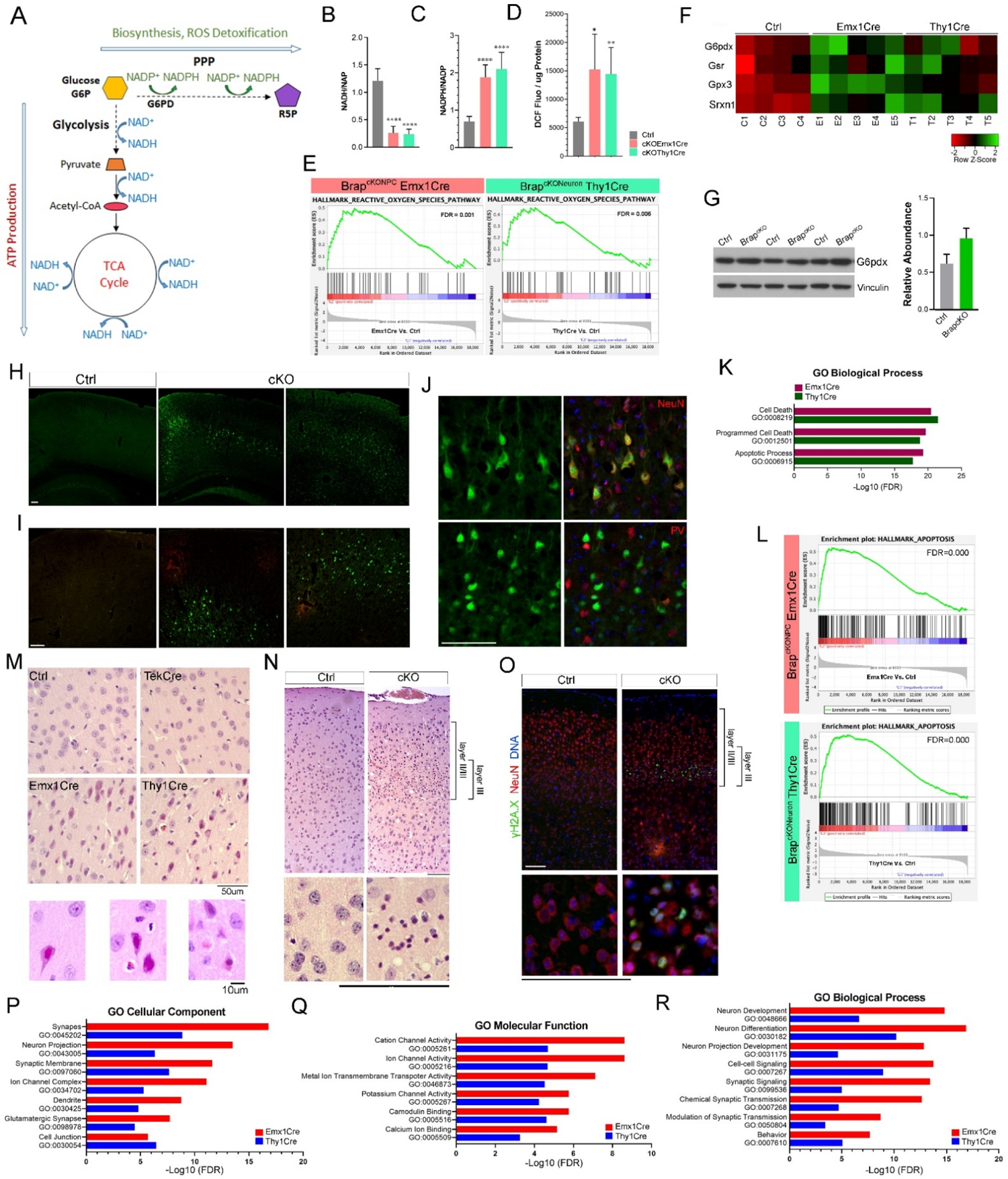
Brain metabolic alterations and neuronal dysfunction in neural BrapcKO mice. (A) A diagram of glucose metabolism and utilization in the brain. (B-D) Relative levels of NADH (measured as NADH/NAD+ ratios), NADPH, (measured as NADPH/NAPD+ ratios), and ROS (measured as DCF fluorescence intensity /ug cortical protein) in cortical tissues of mice with Brap conditional knockout in dorsal telencephalon neural progenitor cells (Emx1Cre), postnatal cortical and hippocampal neurons (Thy1Cre), and Cre-controls. Error bars represent SD; *p < 0.05; ** p< 0.01; *** p< 0.005; **** p< 0.0001 by student’s t-test. (E) GSEA of hallmark pathways reveals the significant enrichment of ROS pathway genes in the cerebral cortical tissue of mice with Brap conditional knockout by Emx1Cre or by Thy1Cre. Shown are enrichment score plots with FDR values compared to cortical genes identified from control mice. (F) Heatmap of expression levels of selected genes associated with oxidative stress in the cortical tissue of mice with Brap conditional knockout by Emx1Cre or by Thy1Cre and aged matched controls. (G) Immunoblotting analysis of G6pdx in cerebral cortical tissues of neural Brap^cKO^ and control mice. (H&I) Representative autofluorescence images of unstained cerebral cortical sections visualized by the green fluorescence channel (H) or both green and red fluorescence channels (I) by fluorescence microscopy. (J) Immunohistological staining of cortical sections with neuronal markers NeuN or parvalbumin PV (both in red) reveals the overlap of green autofluorescent cells with NeuN immunoreactivity in the neocortex of Brap^cKONPC^ mice. Nuclear DNA was stained with Hoechst 33342 in blue. (K) Gene Oncology in Biological Process (BP) terms reveal the enrichments of cell death and apoptosis in upregulated DEGs from mice with Brap conditional knockout by Emx1Cre or Thy1Cre. Shown are selected enrichment terms with corresponding FDR values. (L) GSEA of hallmark pathways reveals the association of apoptosis with unique cortical genes identified from mice with Brap conditional knockout by Emx1Cre or by Thy1Cre. Shown are enrichment score plots with FDR values compared to cortical genes identified from control mice. (M&N) H&E histology images show cortical “red neurons” and “pseudo-laminar necrosis” in neural Brap^cKO^ mice. The position of neocortical layer III is indicated. (O) Representative fluorescence immunohistological images of cortical sections double stained by antibodies to cortical neurons (NeuN, in red) and the DSBs (γH2A.X, in green). Note the high levels of DSBs in layer III cortical neurons of neural Brap^cKO^ mice. Nuclear DNA was stained with Hoechst 33342 in blue. (P-R) Gene Oncology terms identified in significantly down regulated DEGs from cortical tissues of mice with Brap conditional knockout by Emx1Cre or Thy1Cre compared to control mice. Shown are selected enrichment terms in cellular component (CC), molecular function (MF), and biological process (BP), respectively with FDR values. Bars: 100 um or as indicated.

In line with activating PPP towards ROS detoxification, the cortical tissue of neural Brap^cKO^ mice showed significant over-representation of genes in reactive oxygen pathways (Figure 4E). The upregulated transcripts include G6pdx, the rate-limiting enzyme for PPP, Gsr (glutathione reductase), a central enzyme of antioxidant defense, Gpx (glutathione peroxidase), which catalyzes hydrogen peroxide reduction, and Srxn1 (Sulfiredoxin-1), an endogenous antioxidant protein that can prevent oxidative stress (Figure 4F). We further verified increased abundance of G6pdx protein in neural Brap^cKO^ cortices (Figure 4G). As the rate limiting enzyme of PPP, G6pdx determines the reductive power against oxidative damage by producing NAPDH. Consistent with this, we observed strong endogenous fluorescence signals by the microscopic green fluorescence channel in freshly prepared cortical sections of neural Brap^cKO^ mice (Figure 4H-J). Such auto-fluorescence signals match the emission peak of both NADH and NADPH at 440-470 nm^49^. However, NADH was decreased while NADPH and ROS were elevated in neural Brap^cKO^ cortical tissue. Therefore, the green autofluorescence was from NADPH produced by PPP to protect against ROS.

Further supporting the protective role of NADPH in neural Brap^cKO^ mice, we found the cells with green autofluorescence were largely segregated from sites of microbleeds where the autofluorescence from hemoglobin or necrotic cells was shown in both green and red channels (Figure 4I). These green autoflorescent cells were predominantly excitatory neurons, as evidenced by their pyramidal morphology, positive for the pan-neuronal marker NeuN, and rare expression of GABAergic neuronal marker parvalbumin (Figure 4J, Figure S4A). While the attempt of increasing NADPH by turning on PPP is beneficial for fighting against oxidative stress, it further decreases the amount of glucose in glycolytic pathways. Therefore, such glucose metabolic switch may compromise the ATP production, function, and survival of cortical neurons.

As expected, despite the upregulation of antioxidative responses, neural Brap^cKO^ cortices showed significant upregulation of genes associated with cell death and apoptosis (Figure 4K, L), suggesting a dominant impact of metabolic dysfunction on neuronal survival. Although we did not detect apoptotic cells in neural Brap^cKO^ brains by immunoreactivity of cleaved caspase 3, decreases in neuronal density were evident in areas with microbleeds (Figure S4B). We also observed “red neurons” in the cerebral cortex of neural Brap^cKO^ mice (Figure 4M). These acidophilic neurons presented deep red eosinophilic cytoplasm, cell body shrinkage, and pyknotic nuclei, which are characteristic for neuronal damages resulting from hypoxic ischemic brain injuries and usually lead to necrotic neuronal death^50^. In addition, neural Brap^cKO^ mice showed band-like lesions with enrichment of condensed nuclei in neocortical layer III (Figure 4N), which was reminiscent of the pseudo-laminar necrosis resulting from the glucose and oxygen deficiency of cerebral hypoxic-ischemic insults^51^. Notably, DSBs were highly prevalent in these layer III neurons of the neural Brap^cKO^ mice (Figure 4O, Figure S4C). This suggests a specific association between the vulnerability to metabolic stress and DSBs, the feed-forward contribution of vascular defects and ROS to neuronal genome instability, and the synergy between genotoxicity, vascular damage, and oxidative stress in impairing neuronal function and survival.

We further looked into transcripts downregulated in neural Brap^cKO^ cortices to assess neuronal dysfunctions associated with compromised glucose metabolism and increased oxidative stress. Compared with control mice, 1,731 and 757 genes showed significantly decreased expression in cortical tissues of Brap^cKONPC^ and Brap^cKONeuron^ mice, respectively. GO enrichment analyses revealed that these drown-regulated genes are most significantly enriched in neuronal-essential cellular components, such as neuronal projections, dendrites, synapses, and ion channel complexes (Figure 4P). They mainly encode molecules with functions in ion channel activities, metal ion transports, and Ca^2+^ signaling (Figure 4Q), and predominantly engage in biological processes in neuron differentiation, morphogenesis, synaptic signaling, and synaptic transmission (Figure 4R). We also performed GSEA KEGG and reactome pathway analyses, which revealed significant downregulation of genes in long-term potentiation (LTP) in both Brap^cKONPC^ and Brap^cKONeuron^ cortices (Figure S4D). These data together suggest the rapid neuronal function decline of neural Brap^cKO^ mice and strongly support the contribution of vascular and metabolic dysfunction to accelerated neurodegeneration.

## Discussion

Our analysis uncovered a non-cell autonomous effect of neuronal DSBs on cortical glia and vasculature. This finding provides an insight into the synergistic contribution of multiple risk factors for neurodegeneration. While NDs are primarily caused by loss of neurons or neuronal function, they are almost inevitably associated with inflammatory responses of astroglia and microglia as well as alterations in cerebral blood vessels^8–10^. In particular, vascular dementia is the second most common cause after Alzheimer’s disease (AD) for cognitive impairments^52–54^. As the brain uses 20% of the body’s energy to support the constant neuronal activities, it fully relies on uninterrupted blood flow. This requires neurons to make physical and functional interactions with glia and vascular cells, which together form the neurovascular unit (NVU)^55, 56^. Therefore, defects in any constituents of the NVU is expected to result in co-occurrence of neuroinflammation, vascular changes, and neuronal dysfunction. Both cerebral vascular diseases and NDs have a wide range of clinical manifestation and can go silently for decades before causing stroke and dementia at old age. Results of this study highlight an unexpected mechanism connecting neuronal damages with cerebrovascular lesions. The robust cerebral vascular and neurodegenerative phenotype of the neural Brap^cKO^ mouse model also shows a very early onset. It may serve as a new tool for further investigating the etiology, diagnosis, and intervention of both vascular and neurodegenerative dementia.

The main fate of cells with Brap LOF is cellular senescence, driven primarily by recurrent DSBs due to insufficient silencing of satellite repeats^11, 13^. The senescence state permits a direct communication of neurons with their supporting cells. Cellular senescence, though was originally discovered as the state of irreversible cell cycle arrest in culture, presents much more complex phenotypes in vivo. It is a highly dynamic state and can influence multiple other cells with the paracrine effect of SASP. The molecular composition of SASP is context dependent, with which senescent cells can drive normal cells in their neighbor into secondary senescence^57^. We detected the upregulation of a large number of secreted molecules in the cortical tissue of neural Brap^cKO^ mice. Although neuronal DSBs and senescence may primarily account for these molecular changes, secondary senescence of glia and vascular cells or their reaction to senescent neurons likely contribute significantly to the diverse inflammatory and vasoactive molecules that underpin the vascular lesions, neurodegeneration, and mortality of neural Brap^cKO^ mice. Thus, senescence may act not only as a cell autonomous defense response but also a mechanism for promoting synergistic actions in cells of NVU.

Results of this study also reveal a dominant role of neurons and their defects in influencing the cerebral vasculature and glia to drive systemic brain degeneration. Although the molecular exchange between neurons and cerebral blood circulation is highly restricted by the BBB, BBB’s permeability may be increased by SASP-mediated local increase in proteases and inflammatory factors. This suggests a contribution of neuronal senescence to BBB breakdown in AD and other NDs^10^.

Senescence in vivo encompasses cells in heterogeneous states. Besides the association with age-related pathologies, senescence can also be a beneficial process in embryogenesis, wound healing, tumor suppression, and tissue homeostasis^58–60^. Neurons could be considered “senescent cells” by the canonical definition of “irreversible cell cycle arrest”, by the fact that they harbor a large number of somatic mutations^61^, and by their constant secretory features. While this study focuses on the detrimental effect of neuronal DSBs and senescence in inducing cerebrovascular damage, it opens a possibility for future studies to test whether SASP-like secretion may be a physiological mechanism for modulating BBB function and local cerebral blood flow by neuronal activities, as well as whether SASP molecules may serve as therapeutic targets for NDs.

## Materials & Methods

### Mice

Brap floxed (*Brap*^flox/flox^) mice were generated by conventional mouse embryonic stem cell-based gene targeting^62, 63^. Brap^cKONPC^ mice were generated by crossing Brap^flox/flox^ mice with the Emx1-Cre mice purchased from JaxMice (stock # 005628). Brap^cKONeuron^ mice were generated by crossing Brap^flox/flox^ mice with Thy1-cre mice purchased from JaxMice (Stock No: 006143). Brap^cKOEC^ mice were generated by crossing Brap^flox/flox^ mice with the Tek-Cre (Tie2-Cre) mice purchased from JaxMice (stock # 008863). Brap^cKOENeuron^ mice were generated by crossing Brap^flox/flox^ mice with the Camk2a-Cre mice (T29-1) purchased from JaxMice (stock # 005359). All mice used for this study were housed and bred according to the guidelines approved by the IACUC committees of Uniformed Services University of Health Services in compliance with the AAALAC’s guidelines. Experiments were performed using littermates or age and genetic background matched control and mutant groups in both sexes.

### Fluorescence immunohistological and immunochemical analyses

For immunofluorescence staining of mouse cortical tissue, mouse brains were fixed by transcardial perfusion with PBS and 4% paraformaldehyde and then processed in 12um cryosections or 5 um paraffin sections. After treating with antigen unmasking solutions (Vector Labs), brain sections were blocked with 5% goat serum and incubated with primary antibodies in PBS, 0.05% Triton X100, and 5% goat serum at 4°C overnight, and followed by staining with fluorescence conjugated antibodies and Hoechst 33342. Epifluorescence images were acquired with a Leica CTR 5500 fluorescence, DIC, and phase contrast microscope equipped with the Leica DFC7000T digital camera. For immunohistological (IHC) brain analysis, paraffin sections of 5um thickness were treated with antigen unmasking solutions, blocked with 5% goat serum for one hour, incubated with primary antibodies in PBS, 0.05% Triton X100, and 5% goat serum at 4°C overnight. Then the immunosignals were detected by HRP-conjugated secondary antibodies and the DAB substrate using an ABC kit (Vector lab). IHC images were acquired with a Leica CTR 5500 fluorescence, DIC, and phase contrast microscope equipped a Leica DFC7000 T camera. Images were imported to Adobe Photoshop and adjusted for brightness and black values.

### RNA isolation and RT-qPCR

Cerebral cortical tissue was homogenized in TRIzol reagent (Thermo Fisher) followed by total RNA extraction according to the manufacturer’s protocol. 1ug RNA was reverse transcribed into first-strand cDNA using Superscript III reverse transcriptase (Invitrogen). RT-qPCR reactions were performed using Power SYBR Green PCR Master Mix on an Applied Biosystems Step One Plus Real-Time PCR Systems. Primers used for accessing gene expression were synthesized according to validated primer sequences from the MGH-PGA PrimerBank, and are listed in Table S1, RT-qPCR primers. Expression was normalized to TATA-binding protein (Tbp) as an internal control and results were analyzed using the 2^-(ΔΔCT)^ method.

### Bulk RNA sequencing analysis

Purified total RNA from whole cerebral cortical tissue of four control (Brap^flox/wt^ ^Cre-^), six Brap^cKONPC^, five Brap^cKONeuron^, and five Brap^cKOEC^ mice at 3 months of age were processed at the University of Chicago Genomics Facility, where Poly-A+ selected RNA libraries were prepared and sequenced using an Illumina HiSeq 4000 platform (100 bp paired-end reads). RNA sequencing files were transferred to the Tarbell High-performance computing cluster of Center for Research Informatics at the University of Chicago for analysis.

The quality of raw sequencing data was assessed using FastQC v0.11.5. Raw reads were mapped to the mouse reference transcriptome (GRCm38) using Salmon version 1.5.2. After read mapping with Salmon, Bioconductor (https://bioconductor.org/packages/release/bioc/html/tximport.html) was used to read Salmon outputs into the R environment. Annotation data from Gencode vM25 was used to summarize data from transcript-level to gene-level. Filtering was carried out to remove non protein coding or lowly expressed genes. We first removed the non-protein coding genes, which reduced the number of genes from 54418 to 21828. Then genes with less than 1 count per million (CPM) in at least 3 or more samples were filtered out, which reduced the number of genes from 21828 to 14086. A variety of R packages was used for data analysis. All graphics and data wrangling were handled using the tidyverse suite of packages. All packages used are available from the Comprehensive R Archive Network (CRAN), Bioconductor.org, or Github. Differential gene expression analysis was performed using the DESeq2 package. Significance was defined as FDR <0.05 and absolute Fold Change ≥ 1.5.

### Brain metabolic analyses

To assess the glycolic flux and redox status, cortical tissues were dissected from mice that were deeply anesthetized and euthanized by cervical dislocation. The tissue specimens were minced on ice, aliquoted, and stored at −80°C for metabolic analysis. The NADH/NAD+ assay was performed using NAD/NADH Quantitation Colorimetric Kit (BioVision) according to manufacturer’s protocol. The NADPH/NADP+ assay was performed using NADP/NADPH Quantitation Kit (Sigma-Aldrich) according to manufacturer’s protocol. The level of ROS was detected by the OxiSelect™ Green Fluorescence In Vitro ROS/RNS Assay Kit (CELL BIOLABS) according to manufacturer’s protocol. Quantities of cortical proteins were determined by Pierce™ Detergent Compatible Bradford Assay Kit (Thermo Scientific™).

### Evans blue assay

Mice were given an intraperitoneal injection of 2% Evans blue solution (Sigma E2129, dissolved in PBS and filtered through a 0.22um filer, at 4 ml/kg of body weight). The dye was allowed to circulate for 4 hours. Then mice were perfused with 60 ml ice-cold PBS to clear the Evans blue in circulation and euthanized by cervical dislocation. Cerebral cortices were dissected, flash frozen in liquid nitrogen, wrapped in aluminum foil and stored at −80°C. To quantify the amount of Evans blue in cortical tissues, samples were dried in an oven at 55°C, weighted, and immersed in formamide to extract Evans blue from the tissue. After removing tissue precipitates by centrifugation, the amount of Evans blue in the supernatant was determined by absorbance at a wavelength of 620 nm and quantified according to a standard curve. Results were presented as OD 620 per gram of dry cortical tissue.

### Quantification and statistical analysis

No statistical methods were used to predetermine sample size, while all experiments were performed with a minimum of three biological replicates and all cell counts were obtained from at least ten random fields. The experiments were not randomized; the investigators were not blinded to the sample allocation and data acquisition during experiments but were blinded in performing quantitative analyses of immunohistological images using the NIH Image J software. All statistical analyses were done using GraphPad Prism 9.0 software. Data were analyzed by one-way ANOVA or unpaired two-tailed Student’s t tests for comparing differences between different genotypes. Differences were considered significant with a p value < 0.05.

## Supporting information

Table S1 Sequences of RT-qPCR primers

## Acknowledgments

The authors wish to thank the Genomics Facility of University of Chicago for RNA library construction and NGS analysis. This work was supported by startup funds of Uniformed Services University of the Health Sciences to Y.F.

## Author contributions

Y.F. conceptualized the project, designed and performed the experiments, interpreted the results, and wrote the manuscript. C.A.K. performed experiments; C.C.L. initiated the study and performed the experiments; J. Z. performed experiments; W.K. performed data analyses, X.Z. assisted with experiments.

## Disclaimer

The opinions, interpretations, conclusions and recommendation are those of the authors and are not necessarily endorsed by the U.S. Army, Department of Defense, the U.S. Government or the Uniformed Services University of the Health Sciences. The use of trade names does not constitute an official endorsement or approval of the use of reagents or commercial hardware or software. This document may not be cited for purposes of advertisement.

