## Supplemental Figures for "Senescence of cortical neurons following persistent DNA double-strand breaks induces cerebrovascular lesions"

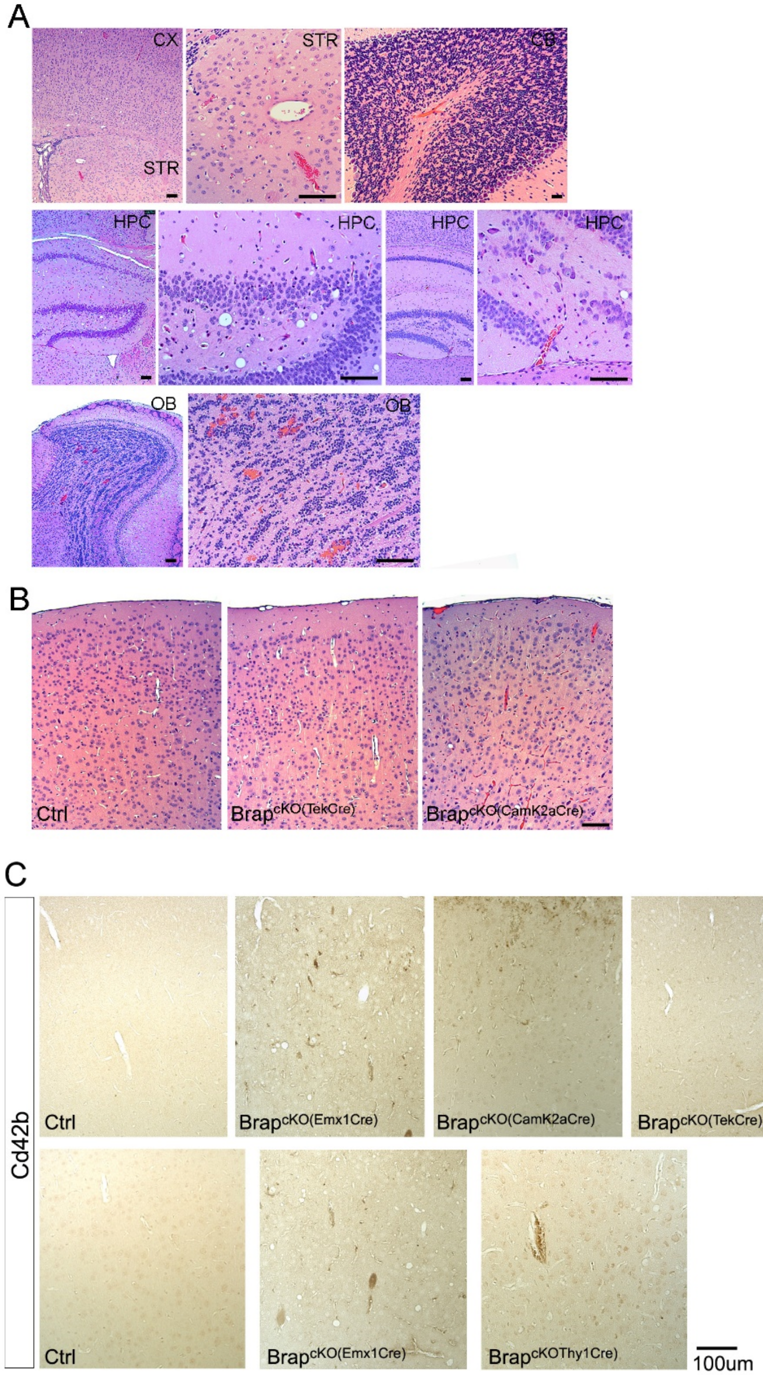


**Figure S1 Brain Histology of BrapcKO mice.**

(A) Representative images of H&E stained neural $Brap^{cKO}$ brains. CX: Cerebral cortex; Str: striatum; CB: cerebellum; HPC: Hippocampus; OB: Olfactory bulb

(B) Representative images of $Brap^{cKOEC}$ and $Brap^{cKOENeuron}$ mice.

(C) Representative Cd42b immunohistochemistry images of various $Brap^{cKO}$ and control mice.

Bars: 100 um

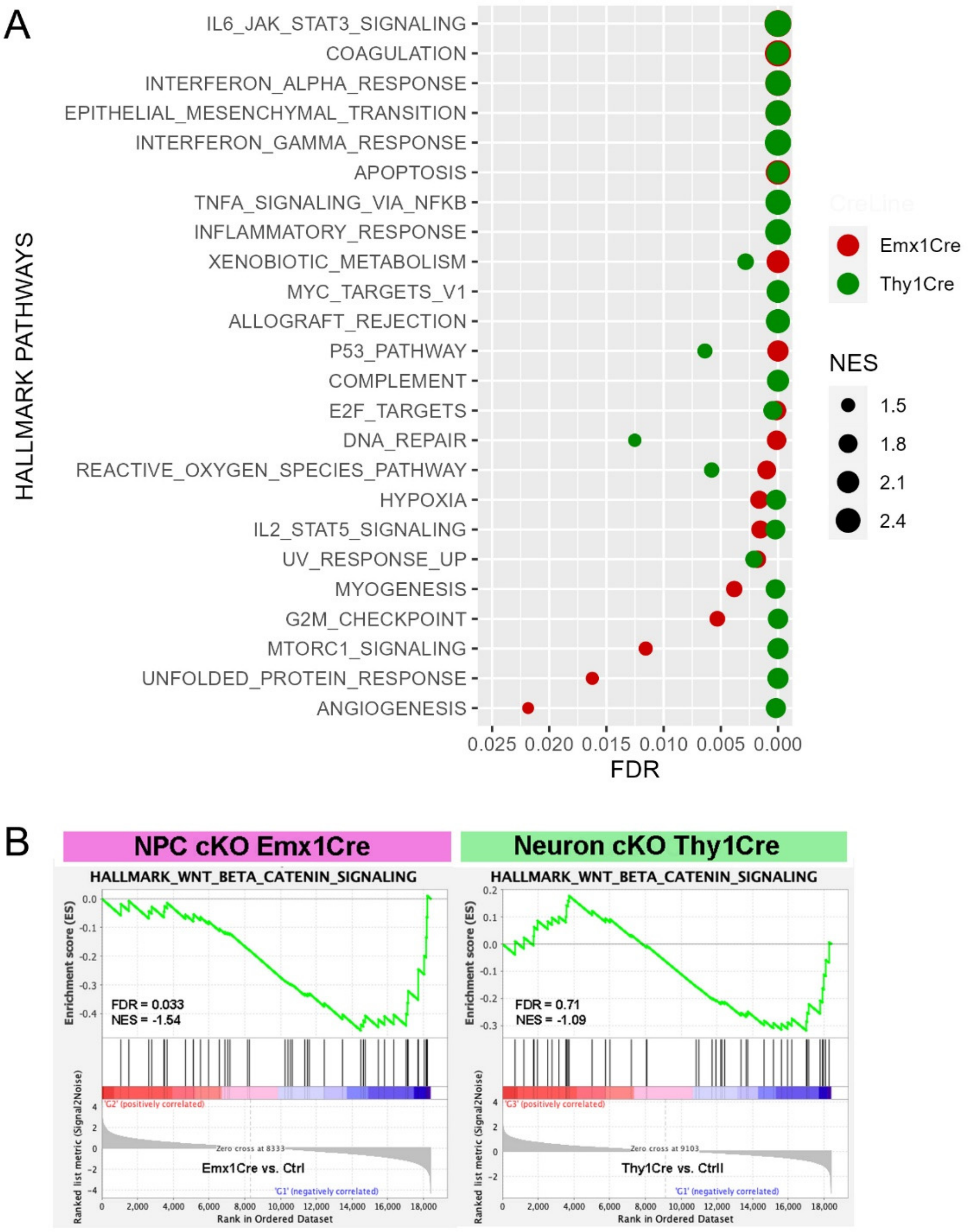


**Figure S2. GSEA of hallmark pathways enriched in cortical genes identified from mice with Brap conditional knockout in dorsal telencephalon neural progenitor cells (Emx1Cre), or postnatal cortical and hippocampal neurons (Thy1Cre).**

(A) Dot plot of hallmark pathways enriched in neural Brap$^{cKO}$ vs control cortical tissues. Shown are FDR values and normalized enrichment scores (NES).

(B) Enrichment score plots with FDR and NES values of hallmark Wnt-Beta Catenin pathway genes enriched in control vs neural Brap$^{cKO}$ cortical tissues.

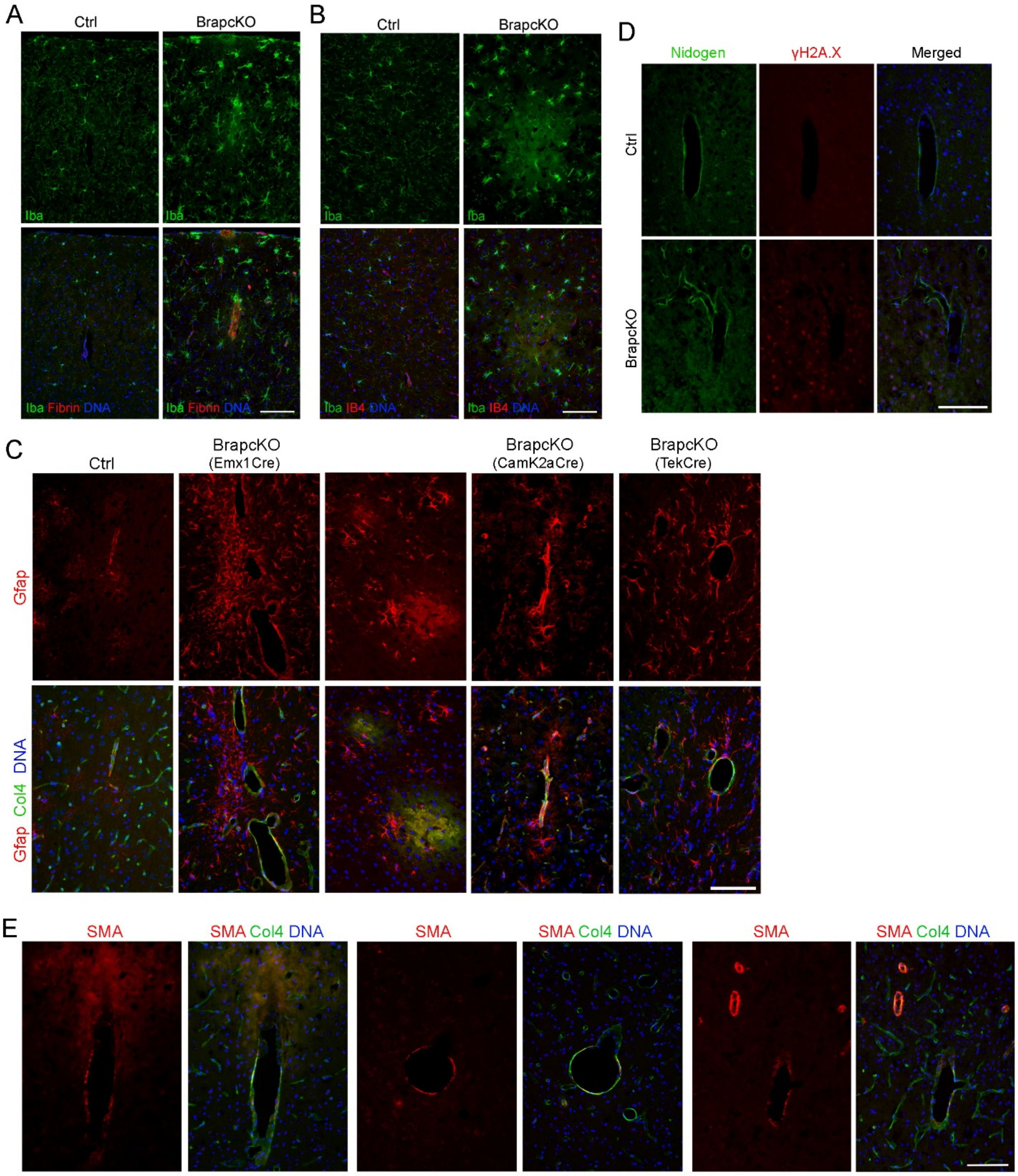

A
Ctrl
BrapcKO
Iba
Iba Fibrin DNA
B
Ctrl
BrapcKO
Iba
Iba IB4 DNA
D
Nidogen
γH2A.X
Merged
Ctrl
BrapcKO
C
Ctrl
BrapcKO (Emx1Cre)
BrapcKO (CamK2aCre)
BrapcKO (TekCre)
Gfap
Gfap Col4 DNA
E
SMA
SMA Col4 DNA

**Figure S3. DNA damage, inflammation and cerebrovascular abnormalities in BrapcKO mice.**

(A) Representative images of cortical sections of Brap$^{cKO}$ mice. Each section was stained by an anti-Fibrinogen (Fibrin, in red) and an anti-Iba antibody (in green).

(B) Representative images of cortical sections of Brap$^{cKO}$ mice. Each section was stained by biotinylated Isolectin 4 (IB4 in red) and an anti-Iba antibody (in green).

(C) Representative images of cortical sections of Brap$^{cKO}$ mice. Each section was stained by an anti-Gfap antibody (in red) and an anti-Col4 antibody (in green).

(D) Representative images of cortical sections of Brap$^{cKO}$ mice. Each section was stained by an anti-γH2A.X antibody (in red) and an anti-Nidogen antibody (in green).

(E) Representative images of cortical sections of Brap$^{cKO}$ mice. Each section was stained by an anti-SMA antibody (in red) and an anti-Col4 antibody (in green).

DNA was stained by Hoechst 33342 in blue. Bars: 100um

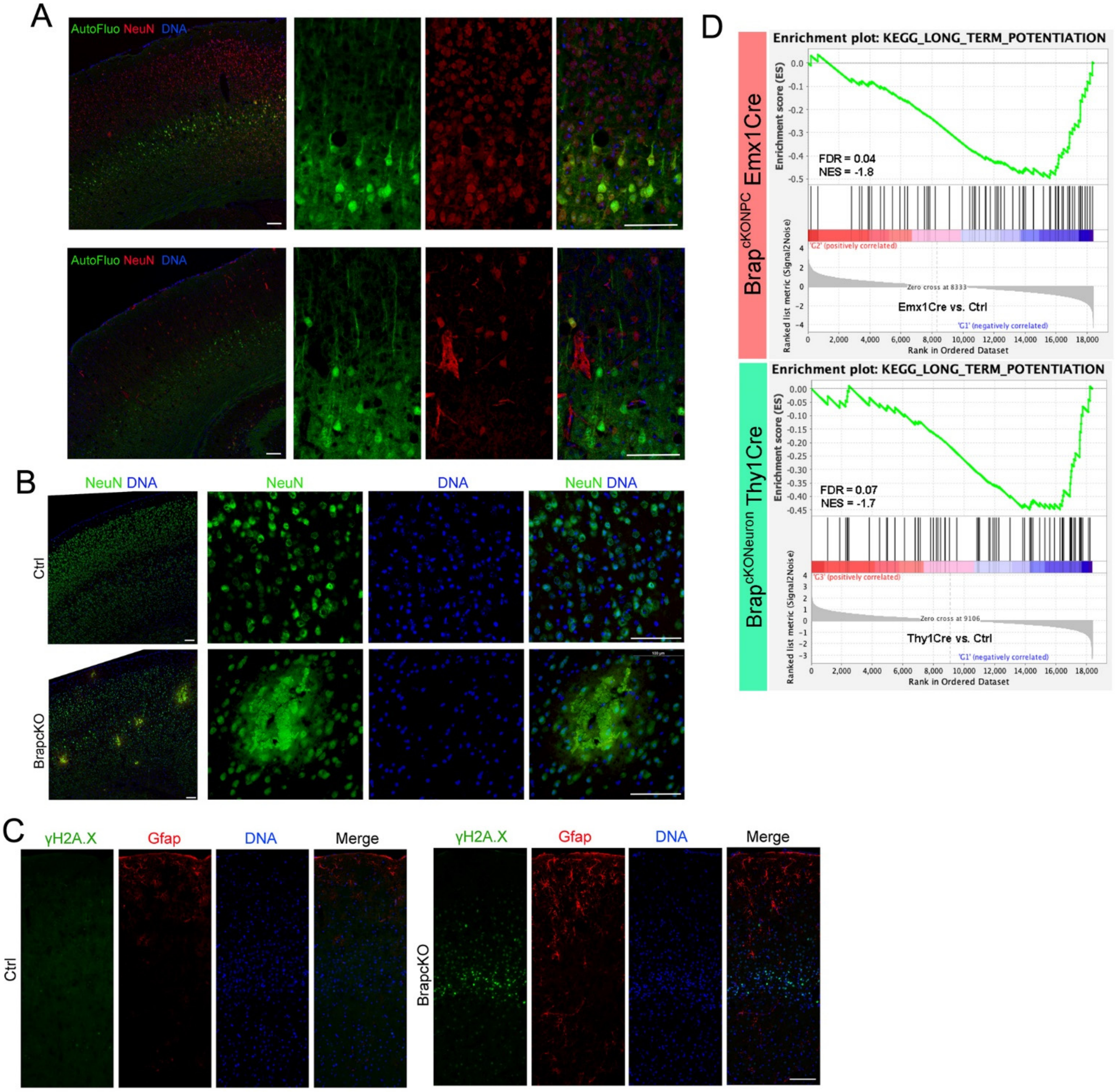
A
AutoFluo NeuN DNA
B
NeuN DNA
NeuN
DNA
NeuN DNA
Ctrl
BrapcKO
C
γH2A.X
Gfap
DNA
Merge
Ctrl
BrapcKO
D
Enrichment plot: KEGG_LONG_TERM_POTENTIATION
Brap cKONPC Emx1Cre
Enrichment score (ES)
FDR = 0.04
NES = -1.8
Ranked list metric (Signal2Noise)
'G2' (positively correlated)
Zero cross at 8333
Emx1Cre vs. Ctrl
'G1' (negatively correlated)
Rank in Ordered Dataset
Enrichment plot: KEGG_LONG_TERM_POTENTIATION
Brap cKONeuron Thy1Cre
FDR = 0.07
NES = -1.7
'G3' (positively correlated)
Zero cross at 9106
Thy1Cre vs. Ctrl
'G1' (negatively correlated)
Rank in Ordered Dataset

**Figure S4. Metabolic stress, necrotic neuronal death, and neuronal function decline in the cerebral cortex of neural Brap$^{cKO}$ mice.**

(A) Representative images of cortical sections of neural Brap$^{cKO}$ mice. Each section was stained by an anti-NeuN or anti-parvalbumin antibody (in red). Immunosignals were merged with autofluorescence and DNA stained by Hoechst 33342 in blue.

(B) Representative images of cortical sections stained by anti-NeuN antibody (in green). Nuclear DNA was stained by Hoechst 33342 in blue. Note the non-specific NeuN immunosignals and reduced cell density in cortical regions with micro-bleeds in neural Brap$^{cKO}$ mice.

(C) Representative images of cortical sections double stained by antibodies against γH2AX (green) and Gfap (red).

(D) Enrichment score plots with FDR and NES values of KEGG pathway of Long-Term Potentiation (LTP) genes enriched in control vs neural Brap$^{cKO}$ cortical tissues.

Bars: 100um.
